# I believe, therefore I am: Effects of self-control beliefs on behavioral and electrophysiological markers of inhibitory and emotional attention control

**DOI:** 10.1101/552570

**Authors:** Naomi Vanlessen, Davide Rigoni, Antonio Schettino, Marcel Brass

## Abstract

In this study, a placebo/nocebo neuro-stimulation procedure was employed to investigate if expectations about self-control can influence self-control exertion. More specifically, we recorded behavioral and electrophysiological responses in an emotional antisaccade task in a between-subjects design, in which one group was led to believe that self-control was enhanced (MSC group) and the other that self-control was weakened (LSC group). This set-up allowed to investigate both response and emotional inhibition, as well as different stages at which control can be exerted during task performance, using Event-Related Potential (ERP) methods. Results showed that the bogus neuro-stimulation indeed installed the expectation of respectively better or worse self-control capacity, as well as the retrospective evaluation at the end of the experiment that the neuro-stimulation changed their self-control in that direction. Participants in the MSC compared to the LSC group showed higher accuracy in trials in which inhibitory control was necessary (antisaccade trials). ERP results showed no effect of the placebo/nocebo manipulation at the level of attention and inhibitory control. In sum, this study showed that high-order cognitive processes are not immune to the influence of expectations induced by a placebo/nocebo procedure, and shows that instructions alone can induce a placebo/nocebo effect in cognitive functioning.

## Introduction

We live in a complex environment that poses countless demands on how we act, regulate our emotions, and comply with complicated rules. The capability to modulate our behaviour and restrain our impulses in order to conform with our own and societal standards is regulated by self-control, the effortful and conscious aspect of self-regulation (Baumeister, Vohs, & Tice, 2007). Self-control can be defined as a set of capacities aimed at changing our way of thinking, feeling, and behaving, in order to regulate urges that could otherwise jeopardize long-term goals, and ultimately well-being (Baumeister, Heatherton, & Tice, 1994). Whether self-control exertion depends on limited resources (Muraven & Baumeister, 2000) or motivational dynamics (Inzlicht, Schmeichel, & Macrae, 2014), previous research has shown that an intact capacity for self-control is essential for the achievement of a wide range of goals, including healthy weight and life style (Hofmann, Adriaanse, Vohs, & Baumeister, 2014; Nederkoorn, Houben, Hofmann, Roefs, & Jansen, 2010), academic and professional accomplishments (Job, Friese, & Bernecker, 2015; Stadler, Aust, Becker, Niepel, & Greiff, 2016), fidelity and wellbeing in intimate relationships (McIntyre, Barlow, & Hayward, 2015; Pronk, Karremans, & Wigboldus, 2011; Visserman, Righetti, Kumashiro, & Van Lange, 2016), and adaptive emotion-regulation (Paschke et al., 2016).

Given the advantages of enhanced self-control and the detrimental consequences associated with its break-down, the present study was aimed at investigating if and how the exertion of self-control can be modulated by beliefs about personal self-control capacity. The study of control-related beliefs has a long-standing tradition in psychology (Ajzen, 2002; Bandura, 1997; Rigoni & Brass, 2014; Rotter, 1966). Recently, researchers started to focus on whether self-control performance is malleable to individuals’ beliefs about their self-control capacity. For instance, in a series of studies Job and colleagues showed that only individuals holding the belief that self-control capacities draw on limited (rather than unlimited) resources showed a decrease in self-control performance after demanding tasks (Job, Dweck, & Walton, 2010; Job, Walton, Bernecker, & Dweck, 2013), attributable to their motivation to take a rest (Job, Bernecker, Miketta, & Friese, 2015), their estimation of how mentally fatigued they were (Clarkson et al., 2016), or their evaluation of resource availability (Clarkson, Hirt, Jia, & Alexander, 2010). In a longitudinal study on students, holding a limited resource theory about self-control was associated with more procrastination, unhealthy eating habits and impulsive money spending, and lower efficiency and grades (Job, Walton, Bernecker, & Dweck, 2015). A limited resource theory is also associated with actual lower self-control and subjective well-being (Bernecker, Herrmann, Brandstätter, & Job, 2015). Similarly, lay theories about self-control influence how many personal goals are strived for by an individual as well as their actual achievement (Mukhopadhyay & Johar, 2005), and the belief that one is able to successfully complete a specific task (i.e., self-efficacy; Bandura, 1977) was found to be predictive of weight loss (Weinberg, Hughes, Critelli, England, & Jackson, 1984) and interest in attractive others when engaged in a long-term relationship (Hamburg & Pronk, 2015). In sum, a wealth of research convincingly shows that beliefs about self-control change outcomes of self-control.

Interestingly, belief changes have also been induced by placebo manipulations. Although focused on similar psychological processes, the belief research is quite detached from research on placebo and nocebo effects. However, the long research tradition in placebo/nocebo treatments in the medical domain demonstrates their potential to induce beliefs about the self. Drugs as well as medical procedures can elicit a placebo/nocebo response and exert both physiological and psychological effects by activating expectancies about outcomes, even though the underlying mechanisms are not fully understood. Although placebo and nocebo effects have been described in a wide range of health-related outcomes including pain analgesia (Caldwell & Gritsenko, 2017) and insomnia (Chung, Sharpe, Glozier, Hackett, & Colagiuri, 2017), their influence on higher-order psychological functions such as self-control has received only little attention. While some recent studies have employed placebo or nocebo instructions to manipulate cognitive processes such as memory (Assefi & Garry, 2003; Parker, Garry, Einstein, & McDaniel, 2011; Sinke, Forkmann, Schmidt, Wiech, & Bingel, 2016; Van Oorsouw & Merckelbach, 2007), performance on knowledge tests (Weger & Loughnan, 2013), and implicit learning (Colagiuri, Livesey, & Harris, 2011), so far only few studies examined placebo effects on cognitive control (e.g., da Gama, Slama, Caspar, Gevers, & Cleeremans, 2013; Schwarz & Büchel, 2015). Nonetheless, the research showing effects of belief manipulations on measures of self-control suggests that this high-level cognitive function is in fact permeable to the influence of suggestions. In the present study, we developed a bogus neuro-stimulation procedure to induce either the expectancy that self-control capacity would be increased in the More Self-Control (MSC) group or decreased in the Less Self-Control (LSC) group. In other words, we capitalized on the placebo/nocebo effects to manipulate people’s belief about their capability to exert self-control.

While placebo/nocebo research has barely treated the topic of self-control at all, in belief research self-control was mainly operationalized in terms of complex outcomes in the social, academic, professional, and health domains. Hence, the precise cognitive and neural underpinnings of belief influences on self-control performance remain unknown. An important aspect of self-control is related to executive functioning, with response inhibition as a crucial feature (Aron, 2007; Inzlicht, Legault, & Teper, 2014). The main goal of the present study was, therefore, to examine whether and how individuals’ beliefs about their own self-control capacity influence self-control exertion in a task targeting inhibition. To this aim, we recorded participants’ electro-encephalogram (EEG) while they completed an emotional antisaccade task (Kissler & Keil, 2008), in which participants are instructed to inhibit a prosaccade – i.e., the automatic orientation of the eyes to a stimulus presented in their visual field – toward neutral or emotionally relevant images. The antisaccade paradigm was designed to measure motor inhibition, as reflexive saccades towards a peripheral stimulus abruptly appearing in the visual field must be suppressed (Hallett, 1978). By using neutral and negatively valenced peripheral stimuli (“targets”), we could also measure whether an emotional target content weakened the capability of suppressing erroneous prosaccades, given the very rapid processing of (negative) emotional information and the increased attentional capture by such stimuli (Pourtois, Schettino, & Vuilleumier, 2013). Hence, by combining the usage of an emotional antisaccade task and EEG recordings, we could investigate whether both behavioral and emotional inhibition were both impacted by the placebo/nocebo neuro-stimulation. Crucially, the high temporal resolution of EEG also allowed us to pinpoint the exact locus of impact of placebo/nocebo-induced beliefs about self-control. More precisely, by studying different Event-Related Potentials (ERPs) reflecting specific cognitive processes, we could distinguish between the impact of our manipulation on anticipatory proactive control and reactive control processes (Braver, 2012). With regards to proactive control, we looked at the negativity ramping up between the cue and the target, known as the Contingent Negative Variation (CNV; Walter, Cooper, Aldridge, McCallum, & Winter, 1964). The CNV reflects preparatory control and is typically larger in conditions demanding more control (antisaccade trials) compared to less control-demanding conditions (prosaccade trials; Ansari & Derakshan, 2011; Klein, Heinks, Andresen, Berg, & Moritz, 2000; Richards, 2013). Additionally, we investigated the impact of the placebo/nocebo suggestions on attentional processes, in line with recent research showing proactive modulations of attentional processes in task contexts that demand the inhibition of a target in a proportion of trials (Langford, Krebs, Talsma, Woldorff, & Boehler, 2016). To that aim, we measured the P1 component following target onset, reflecting gain control of selective attention in extrastriate visual areas (Hillyard, Vogel, & Luck, 1998; Luck, Woodman, & Vogel, 2000) and sensitive to the emotional valence of stimulus material (see Pourtois et al., 2013 for a review). To quantify reactive control processes, we investigated the N2/P3-complex, considered a physiological marker of inhibition. Post-target activity in the antisaccade task typically shows the pattern of an enhanced N2 and a decreased P3 in anti-compared to prosaccades, (Mueller, Swainson, & Jackson, 2009; Vanlessen, De Raedt, Mueller, Rossi, & Pourtois, 2015).

Based on the characteristics of the antisaccade paradigm and these ERPs, we formulated the following hypotheses. At the behavioral level, we expected that the LSC group would show a lower accuracy in trials necessitating inhibitory control (i.e., the antisaccade trials) compared to the MSC group. At the electrophysiological level, we hypothesized that the activity preceding targets would be lower in the LSC compared to the MSC group for antisaccade trials at the level of the CNV, indicative of a dampened proactive preparation in this group, in addition to the classical stronger preparatory activity in anti-compared to prosaccade trials. Regarding the N2/P3 complex, we expected that the classical increased N2 and decreased P3 in pro-compared to antisaccade trials would be more pronounced in the MSC group. If the placebo/nocebo manipulation also impacted processing at the attentional level, we would also see a group difference on the visual P1, with lower amplitudes in the MSC compared to the LSC group, especially for emotional images.

## Methods

### Participants

Eighty-two healthy volunteers participated after giving written informed consent^1^. Two participants did not complete the experiment because of technical issues. Data of 14 participants were excluded from further analysis because the manipulation check indicated that the self-control manipulation was not successful (i.e., they did not report to expect the predicted change in self-control due to placebo brain stimulation) and of two other participants because they did not execute the task correctly. The remaining participant sample consisted of 64 participants (15 males) with a mean age of 21 years (SD = 3.20) ranging between 17 and 42 years, with 32 participants per group. The study was approved by the local ethics committee and in accordance with the Declaration of Helsinki.

### Materials

Placebo neuro-stimulation (i.e., tDCS) was used to induce beliefs about the efficiency of brain mechanisms underlying intentional behavior. Prior to participation, volunteers received an information folder by email to explain the functioning of tDCS, the procedure of the study, and selection criteria. Crucially, they were informed that their prefrontal cortex would be stimulated with the aim to influence self-control processes (da Gama et al., 2013). One group of participants was led to believe that their ability to intentionally exert self-control would be enhanced (More Self-Control group, MSC), while the other group was informed that their intentional self-control capacities would be weakened (Less Self-Control group, LSC)^2^. Prior to participation, volunteers were required to fill out a safety checklist for tDCS to strengthen the believability of the bogus neurostimulation. To avoid that participants would find it suspicious that they were informed about the expected effect of the stimulation, they were told a cover story stating that the nature of the study obliged the researchers to explain the procedure in detail prior to participation. For the actual bogus neurostimulation, a round trembling device (~1 cm diameter) was placed over the right prefrontal cortex and was activated for 5 minutes, in order to give the impression of active offline neurostimulation. A real tDCS device was presented to the participants and programmed for the stimulation in their presence. The trembling started following two short beeps and ended after another short beep.

In the antisaccade task, participants were instructed to reorient their gaze from a white 1.7° x 1.7° fixation cross in the middle of the screen either towards (in prosaccades) or away from (in antisaccades) a target appearing right or left from fixation. The fixation cross turned either green or red, indicating whether a prosaccade or an antisaccade had to be executed, respectively. Participants were instructed to fixate the target for its entire presentation. In 50% of the trials, the targets consisted of neutral images; in the other 50% the targets were arousing images with a negative valence. In total, 80 neutral and 80 negative targets were presented (see Fig. 1C for an example). The images covered a rectangle of 10.7° x 8.4° visual degrees on the right or left side of the fixation cross, on the same horizontal axis at 3.5° distance from the center of fixation. Each block consisted of an equal number of pro- and antisaccade trials, with half of the targets occurring on the left and half on the right side of the fixation cross in random order, half of which were neutral and half were negative. No fixation cross was presented during target presentation. All stimuli were presented against a uniform black background. The task was programmed using E-Prime Version 2 (Psychology Software Tools, Inc., 2001).

**Figure 1.**
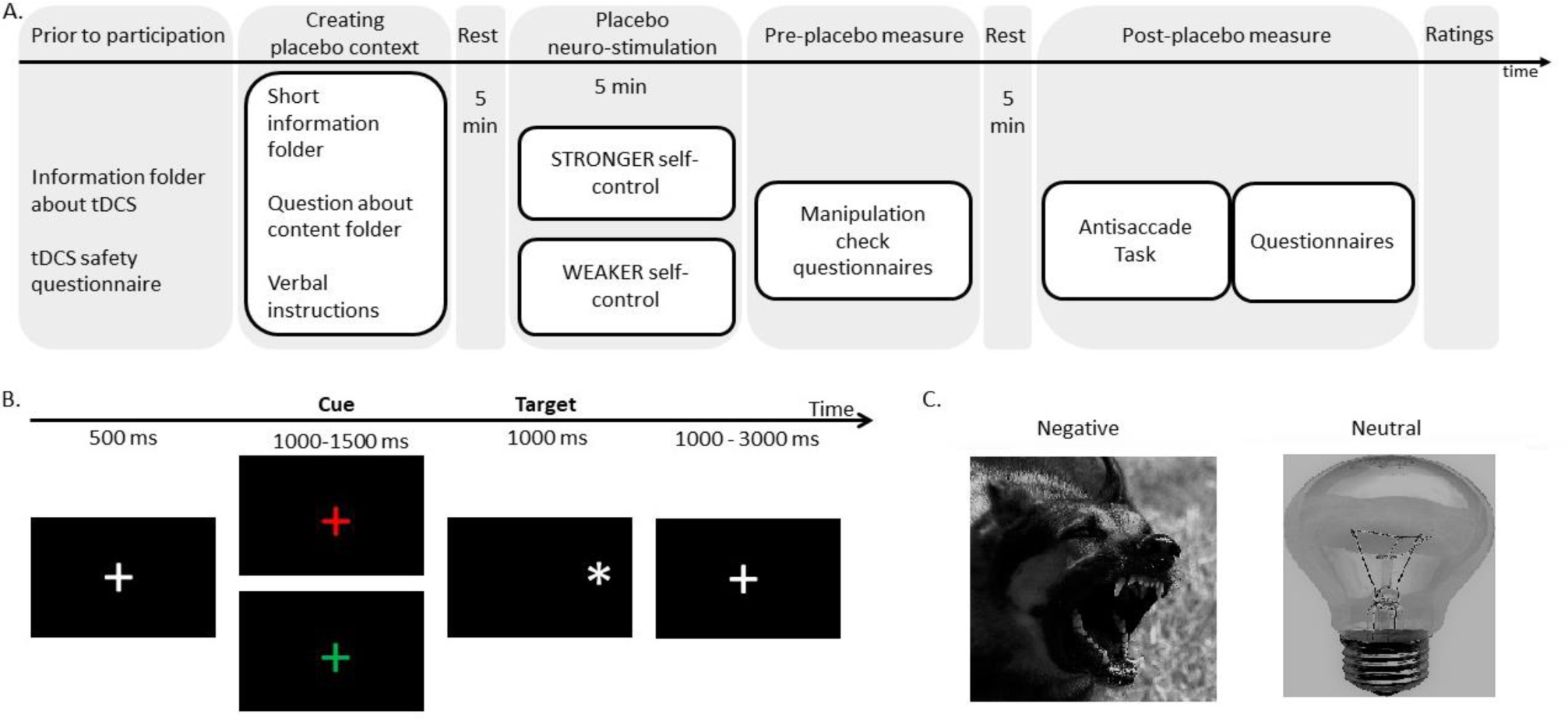
(A) Procedure and (B) trial sequence. The fixation cross turning green instructed participants to make a saccade towards the image (prosaccade), while a red fixation cross instructed them to make a saccade away from the image antisaccade); each type of saccade occurred in 50% of the trials. Note that in the actual experiment, no white star was shown but an image as can be seen in (C). Example of a neutral and a negative image that was used as a target in the antisaccade task. Note that these pictures are examples taken from Wikipedia Commons.

### Images

The images were collected from the Nencki Affective Picture System (NAPS; Marchewka, Żurawski, Jednoróg, & Grabowska, 2014), the International Affective Picture System (IAPS; Lang, Bradley, & Cuthbert, 2008), the Geneva Affective Picture Database (GAPED; Dan-Glauser & Scherer, 2011), and the Emotional Picture System (EmoPicS; Wessa et al., 2010). All images were gray-scaled to exclude the influence of possible color differences between negative and neutral stimuli on visual processing. The negative and the neutral images contained an equal number of images depicting living (41) and non-living (39) objects each. Taking both principal emotion dimensions into account (Russell, 1979), the used negative images have a lower mean valence compared to neutral images (negative: *M* = 3.3, *SD* = 0.4; neutral: *M* = 5.3, *SD* = 0.5; *Q* = 592.10, *p* < .001), as well as a higher arousal level (negative: *M* = 5.6, *SD* = 0.5; neutral: *M* = 4.1, *SD* = 0.7; *t* = 129.41, *p* < .001), based on the scaling of the normalization of each respective scale. Additionally, apparent contrast and measures of complexity (JPEG size and sub-band entropy, Rozenholtz, Li, & Nakano, 2007) were not different between neutral and negative images (*Q* < 0.4, *p* >.5).

At the end of the main task, image ratings were required to verify that participants perceived valence and arousal of the picture set in accordance with the normative ratings. The 80 neutral and 80 negative images from the antisaccade task were presented in random order in four blocks (20 of each valence per block; each block contained the same stimuli as the corresponding task block). In addition, each image was presented for the same duration as in the antisaccade task (i.e., 1000 ms), in order to approach the task situation during the rating as much as possible. Each image was followed by a Self-Assessment Scale (SAM) for arousal and a SAM for valence (Bradley & Lang, 1994). Participants assigned a score ranging from 1 to 9 on each scale using the buttons on the numeric pad.

### Procedure

The procedure consists of several phases, including the placebo/nocebo neuro-stimulation, task execution, and questionnaires and picture ratings at the end of the experiment (see Fig. 1A).

Upon subscription to the experiment via an online platform (Experimetrix), participants received an information folder and a tDCS safety checklist via email (similar to da Gama et al., 2013), to induce the expectancy of real brain stimulation and its capacity to change self-control exertion. Upon arrival in the lab, participants were seated in front of a computer screen, filled out a paper version of the tDCS safety checklist, and were given an information folder containing specific information about the experiment (i.e., whether to expect self-control to increase or decrease). The experimenter then shortly repeated the procedure and what effect participants could expect on their self-control capacity, after which participants were asked to write down how they expected that the brain stimulation would influence their self-control. This procedure allowed us to check whether participants read the folder and understood the expectancies we wanted them to hold regarding the influence of the tDCS bogus stimulation. In case participants did not correctly recapitulate the expected influence of tDCS, they were invited to read the folder again. Finally, participants signed the informed consent and were asked to report their age and handedness.

After this belief induction phase, participants were prepared for EEG recording followed by 5 minutes of rest with eyes closed (data will not be discussed here). Next, participants were asked to remained quiet for five minutes with eyes closed (data will not be discussed here). The bogus tDCS electrode (i.e., trembling element) was then placed on the head of the participant. Next, participants completed questions about their expectancies of the brain stimulation effects on their task performance (see Appendix 1 for a list of the questions and response options). Then, the bogus tDCS stimulation was activated during five minutes. Participants were seated and instructed to remain still during this time.

The belief induction phase was followed by five minutes of rest with eyes closed (data will not be discussed here). Next, four practice trials followed by four blocks of 40 trials of the antisaccade task started, separated by a self-paced break. Each trial started with the presentation of a fixation cross (500 ms), followed by a cue (random duration between 1000 and 1500 ms), a target (1000 ms), and again a fixation cross (random duration between 1000 and 3000 ms; see Fig. 1B).

After the task, participants completed questions about the influence of the bogus brain stimulation on task performance (see Appendix 1), as well as the General Self-Efficacy Scale (GSE; Schwarzer & Jerusalem, 1995), the Multidimensional Iowa Suggestibility Scale (MISS; Kotov, Bellman, & Watson, 2004), and the Free Will Inventory (FWI, Nadelhoffer, Shepard, Nahmias, Sripada, & Ross, 2014). Finally, participants completed the picture ratings (see above).

### Acquisition and reduction of EEG data

Continuous EEG signal was recorded with 64 Ag/AgCl electrodes using Biosemi Active Two System, referenced online to the CMS-DRL ground, and digitized at 512 Hz. Offline, data were further processed and analyzed using Brain Vision Analyzer (Brain Products GmbH, Munich, Germany). Scalp EEG signal was referenced to the linked mastoids and a 0.016 Hz high-pass filter, a 70Hz low-pass filter, and a 50 Hz Notch filter were applied. Noisy channels were interpolated using a spherical spline procedure (Perrin, Pernier, Bertrand, & Echallier, 1989), resulting in a mean of .98 (SD = 1.30, range from 0 to 5) interpolated channels per participant, with no difference between groups (*t*(62) = .49, *p* = .63). Next, the data was segmented in large epochs (−300 to 4000 ms cue-locked) to define the accuracy of eye movements in each trial, after which ocular correction was applied using the Gratton algorithm to automatically correct artifacts caused by eye movements (Gratton, Coles, & Donchin, 1983). Next, the signal was baseline-corrected using the 300 ms prior to the cue. The large epochs of correct trials only were then segmented in target-locked epochs for analysis of activity preceding (−1000 to 600 ms) and following (−300 to 1000 ms). These epochs were again baseline-corrected (for pre-target activity: from 1000ms to 900 ms before target presentation; for post-target activity: the 300 ms preceding target presentation) and artifacts were rejected semi-automatically using an absolute voltage criterion of ±100 μV. With this criterion, 14,53 % of the total epochs were rejected.

Individual averages were calculated separately for the pre- and post-target components, and separately for the pro- and antisaccade trials. For the post-target activity, averages were also made separately per emotion condition (neutral vs. negative). For the latter, we used surface Laplacians estimates from the averaged individual monopolar EEG signal for the CNV increasing prior to target onset of the last 400 ms prior to target-onset. Spherical spline interpolation procedure was applied and the second derivatives in the two dimensions of space were computed (degree of spline = 3, maximum degrees of the Legendre polynomial = 15). Given that the Laplacian improves the spatial resolution of the signal, the CNV was expected to be maximum over fronto-central and central electrodes. Therefore, we FCz and Cz were used to calculate the mean CNV activation. For post-target activity, we measured the average activity from 80 to 90 ms for the P1, on the electrodes where this peak was maximal bilaterally (PO7, PO3, O1, PO8, PO4, O2). We performed a peak analysis^3^ for the N2 (between 180 and 260 ms) on fronto-central electrodes (Fz, F1, F2, AFz and FCz) selected based on the topographical properties of this component for each average individually (i.e., right/left; negative/neutral; pro/anti). Next, average amplitude of 20ms around the peak was calculated. The topography of P3 showed a bilateral activation in posterior regions. In line with this topography, we measured the mean activity between 250 and 400 ms post-target onset on eight occipital electrodes (P5, P3, PO7, PO3, P8, P6, PO4 and PO8). Activity for targets presented right and left was averaged together given that we had no hypotheses about an influence of presentation location on any of the dependent variables. For the post-target ERPs, we submitted the mean amplitude values of all electrodes to a mixed model ANOVA with saccade and emotion as within-subject factors and group as between-subject factor.

### Analysis

All statistical analyses were performed using JASP. For the manipulation check, t-tests were performed on VAS scores to the question before and after the task to check if participants believed the bogus neuro-stimulation had an influence on their task performance (corrected for multiple comparisons). T-tests were also used for checking differences on the questionnaire scores (corrected for multiple comparisons). Descriptive statistics were checked for both groups together for the other questions, as they served to verify that participants believed the manipulation, but no difference between groups was expected.

Repeated measures ANOVAs with within-subject factors Saccade (pro vs. anti) and Emotion (neutral vs. negative), and between-subject factor Group (MSC vs. LSC) were performed on the mean accuracy rates, and the pooled means for the P1, N2, and P3 separately. A repeated measures ANOVA with within-subject factors Saccade (pro vs. anti), and between-subject factor Group (MSC vs. LSC) was performed on the pooled mean for the CNV. Significant interactions were followed-up by t-tests. For accuracy and the CNV, an additional explorative ANOVA was performed including Half (first half vs. second half of the experiment) as an extra within-subject factor.

For the picture ratings, the mean SAM scores for valence and arousal were compared in separate ANOVAs with Emotion as within-subjects factor to check that neutral and negative stimuli were indeed rated differently.

## Results

### Behavioral results

#### Manipulation check and questionnaires

Prior to the bogus brain stimulation, participants in the MSC group indicated to expect that their level of self-control would be increased (M = 2.16, SD = 1.87) and participants in the LSC group reported to expect decreased levels of self-control (*M* = −2.26; SD = 1.76, t(61) = 9.67, p <.01, d = 2.44). Similarly, after they completed the task, participants in the MSC group indicated to believe that the brain stimulation had enhanced their performance (*M* = 2.04, SD = 1.45) compared to participants in the LSC group that indicated that it decreased their performance (M = −2.21; SD = 1.68, t(62) = 10.66, p <.01, d = 2.72; see Fig. 2A).

**Figure 2.**
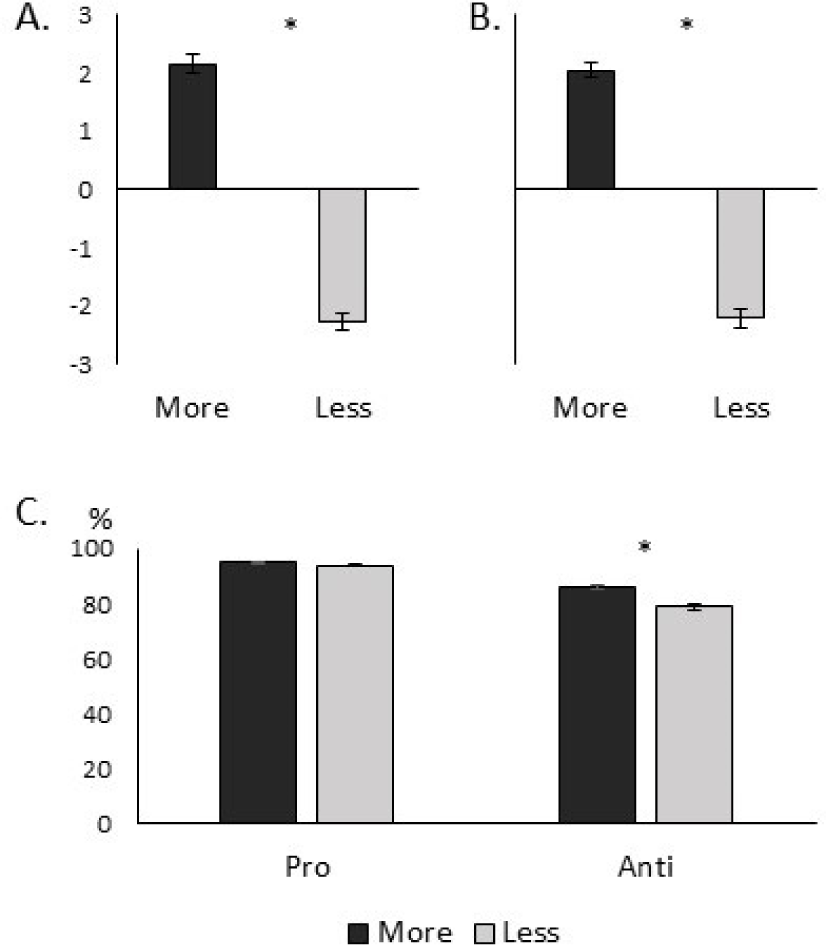
Results of the manipulation check, showing participants in the MSC group (More) expected that they will experience better self-control during the task compared to LSC group (Less) before the task (A) and estimated that their self-control was bettered during the task at the end of the experiment (B), while the LSC group estimated their self-control was attenuated during task performance. (C) Behavioral accuracy. More = More self-control (MSC); Less = Less self-control (LSC).

Furthermore, participants showed that they were confident about the efficacy of the (bogus) brain stimulation to modulate their self-control capacity (see Table 1), indicating that they believed the stimulation would be successful in eliciting the expected result (tDCS expectancy; *M* = 5.17, SD = 0.92), and that both on a subjective (feel; *M* = 4.17, SD = 1.32) and a rational level (think; *M* = 4.78, SD = 0.95) that it would change the performance of the task at the beginning of the experiment. Participants in both groups indicated that at the end of the experiment they still believed that the brain stimulation influenced task execution (tDCS estimation; *M* = 4,36, SD = 1.26) and that they would attribute good/bad performance to the bogus stimulation (attribution; *M* = 4.16, SD = 1.33). Furthermore, participants indicated to find the antisaccade trials quite effortful (effort; *M* = 4.70, SD = 1.19) as well as that they felt capable of executing the task well (task efficiency; *M* = 4.56, SD = 1.10).

Finally, the SEQ, MISS and FW questionnaires did not show any statistically significant differences between groups (all *p*s > .05).

#### Ratings images

Participants rated the unpleasant images as more arousing (M = 4.31, SD = 1.53; t(63) = 10.22, p < .01, d = 1.23) and more negative (M = 3.38, SD = 0.60; t(63) = 20.16, p < .01, d = 2.52) compared to neutral images (Arousal: M = 3.05, SD = 1.36; Valence: M = 5.17, SD = 0.42), confirming the emotion differences between picture classes suggested by the normative ratings.

#### Accuracy

The omnibus ANOVA showed that participants scored higher on prosaccades compared to antisaccades (F(62) = 73.61, p < .01, ω^2^ = .31), as well as an interaction effect between trial and group (F(62) = 4.26, p < .04, ω^2^ = .02). No other main or interaction effects reached statistical significance (p > .36). Follow-up t-tests showed that accuracy was significantly lower in the LSC (M = 78.94%, SD = 14.03) compared to the MSC (M = 85.86%, SD = 10.33; t(62) = 2.25, p = .03, *d* = 0.56) in the antisaccade trials. No significant group difference was found in the prosaccades (MSC: M = 95.07%, SD = 3.36; LSC: M = 93.99%, SD = 4.40; t(62) = 1.11, p = .27, *d* = 0.28; see Fig. 2B).

### ERP results

#### CNV

The ANOVA revealed a main effect of Saccade (*F*(1,62) = 4.53, *p* = .04, ω^2^ = 0.02), with the CNV being larger in the antisaccade (*M* = −0.28 µV/cm^2^, *SD* = 0.35 µV/cm^2^) than in the prosaccade trials (*M* = −.19 µV/cm^2^, *SD* = .24 µV/cm^2^; see Fig. 3A). Neither the Saccade x Group interaction (*F*(1,62) = 0.69, *p* = .41, ω^2^ < .01) nor the main effect of Group (*F*(1,62) = 2.82, *p* = .098, ω^2^ = .03) reached significance.

**Figure 3.**
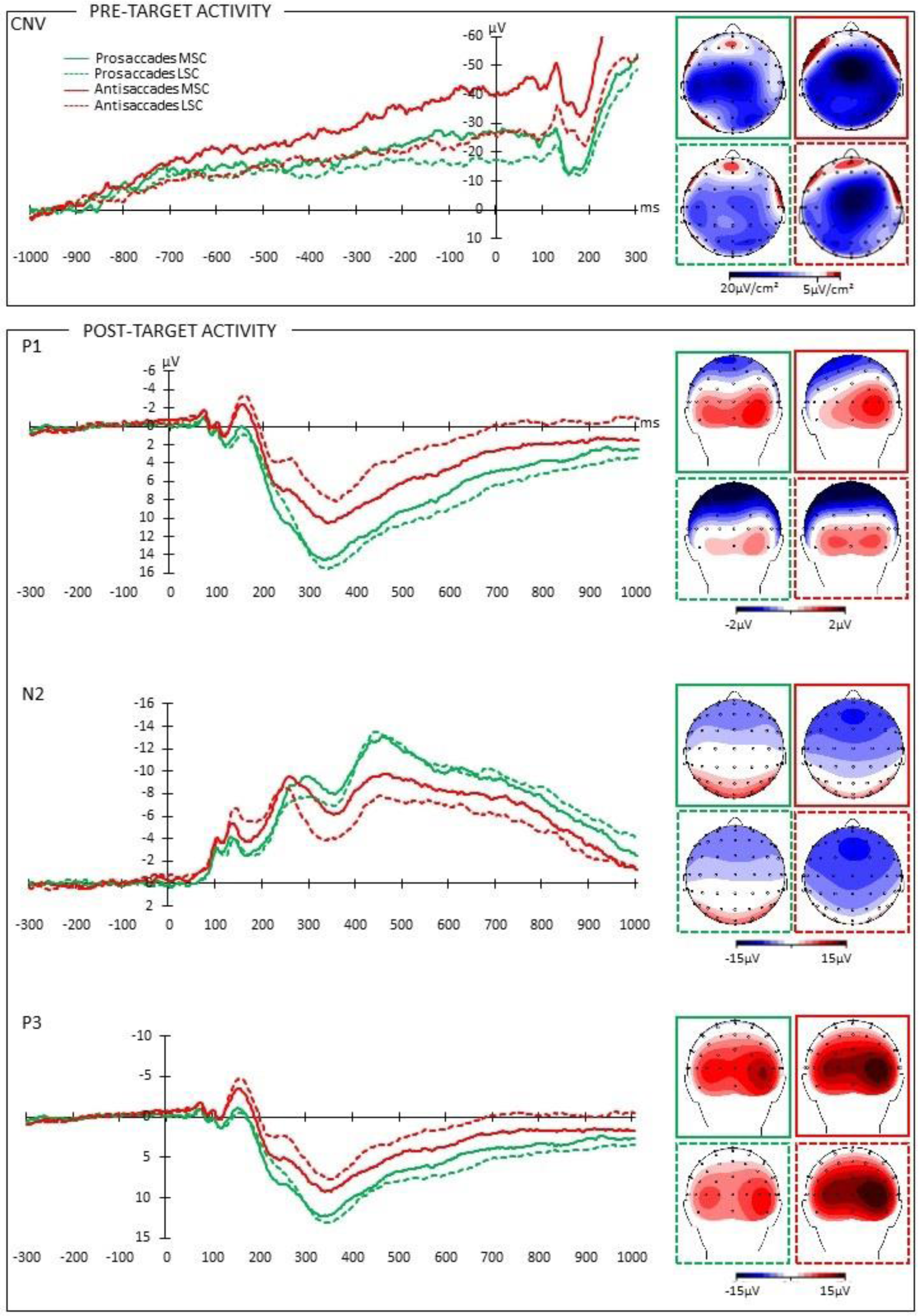
Grand average ERPs for the CNV, P1, N2, and P3 components, pooled over the considered electrodes (FCz and Cz for the CNV; PO7, PO3, O1, PO8, PO4, O2 for the P1; Fz, F1, F2, AFz, FCz for the N2; P5, P3, PO7, PO3, P8, P6, PO4, PO8 for the P3; FCz, Cz for the CNV) and the corresponding voltage maps. MSC = More self-control group; LSC = Less self-control group. The 0 on the horizontal axes represents target onset. Negative values are plotted upwards.

#### P1

The amplitude of the P1 was significantly higher in the prosaccade trials (M = 0.66, SD = 2.09) compared to the antisaccade trials (M = −0.17, SD = 2.02; F(1,62) = 10.48, p = .002, ω^2^ = .04), as well as for neutral (M = 0.50, SD = 1.77) compared to negative images (M = −.004, SD = 2.10; F(1,62) = 6.01, p = .02, ω^2^ = .01). We also found a trial x emotion interaction (F(1,62) = 4.49, p = .04, *ω^2^* = .01; see Fig. 3B), due to a significant difference between neutral (M = 1.13, SD = 2.24) and negative images (M = 0.20, SD = 2.60) in prosaccade trials (t(63) = 3.02, p = .004) but not in antisaccade trials (neutral: M = −0.14, SD = 2.20; negative: M = −.21, SD = 2.35; t(63) = 0.27, p = .79). Group did not influence the P1 amplitude (all p > .05), although a trend was observed in the trial x group interaction (*t*(62) = 3.70, *p* = .059, *ω*^2^ = .01).

#### N2/P3

The N2 was larger in amplitude in antisaccade trials (M = −10.76, SD = 4.63) compared to prosaccade trials (M = −8.50, p = 4.04; F(62) = 27.86, p < .01, ω^2^ = .06; see Fig. 3C). No other effects reached significance (all *p*s > .13). The amplitude of the P3 was larger in prosaccade trials (M = 9.98, p = 4.15) compared to antisaccade trials (M = 4.24, SD = 4.10; F(62) = 266.34, p < .01, *ω*^2^ = .33). No other significant effects were present (all *p*s < .22).

### Exploratory analysis

To check for the possibility the belief effects faded out over time, we also checked if a stronger effect of the manipulation was present in the first compared to the second half of the experiment in both the accuracy and the CNV. For the behavioral results, this analysis showed indeed a trial x half x group effect (*F*(62) = 4.25, *p* = .04, *ω*^2^ = .008), which was driven by the trial x group interaction in the second half only (*F*(62) = 5.69, *p* = .02, *ω*^2^ = .03) while no such interaction was present in the first half (*F*(62) = 0.95, *p* = .34, *ω*^2^ < .01). For the CNV, no three-way interaction was found (*p* = .68).

## Discussion

Due to its paramount importance for interpersonal, academic, and professional success, self-control or the ability to inhibit impulses to align behavior with rules and long-term goals, is a long-time subject of philosophical and psychological inquiry. One area of research has specifically focused on the modulatory power of beliefs and convictions to enhance or weaken self-control (Ajzen, 2002; Bandura, 1997; Rigoni & Brass, 2014; Rotter, 1966). In the current study, we build on this tradition to investigate if and how changed beliefs about self-control, induced using placebo/nocebo neuro-stimulation, can modulate self-control performance on a behavioral and electrophysiological level in an antisaccade task. More precisely, we investigated its effect on behavioral response inhibition, and on attention-(P1) and inhibition-related ERPs (N2/P3-complex), as well as preparatory activity (CNV).

Self-reports indicated that the placebo/nocebo-inducing instructions elicited the intended expectation about self-control exertion prior to the experiment, and that participants also retrospectively judged their performance to be influenced by the neuro-stimulation. Especially the latter suggests that the participants’ belief that their self-control was enhanced/weakened by the bogus tDCS was active during task performance. The placebo/nocebo tDCS impacted behavioral performance (and was associated with a numerical difference in P1 and CNV amplitude) in a trial-specific manner, showing that the beliefs specifically impacted the trials in which control was actually necessary (antisaccade trials), but not when very low levels of control sufficed to perform the task (prosaccade trials). While accuracy in the prosaccade trials did not differ between the two groups, participants in the LSC group obtained lower accuracy compared to the MSC group in the antisaccade trials. Groups did not influence post-target processing differentially at the attentional level, nor at the level of the later inhibition-related N2/P3-complex. Trait questionnaires showed that groups did not differ in suggestibility, self-efficacy, or the extent they supported the notion of having free will. The results and their implications are discussed below.

### Placebo/nocebo and self-control

A long tradition in clinical research has described placebo and nocebo effects in a wide range of health-related outcomes, and recently have also been observed in emotion and pain regulation (Petrovic et al., 2005; Wager et al., 2004), and cognitive processes (Assefi & Garry, 2003; Colagiuri et al., 2011; Parker et al., 2011; Sinke et al., 2016; Van Oorsouw & Merckelbach, 2007; Weger & Loughnan, 2013). One largely overlooked area in this research is the influence of placebo and nocebo on higher-order psychological functions engaged in complex behavior, such as self-control. However, extensive belief research suggests that this high-level cognitive function is in fact malleable to suggestions. One placebo study employed sham neuro-stimulation to induce the belief that colour perception would either be enhanced or weakened (da Gama et al., 2013). After the bogus neuro-stimulation, participants completed a colour Stroop task, in which they had to name a colour word (e.g., the word “yellow”) displayed in another colour (e.g., red font). Better (lower) accuracy was found in the group of participants believing that the device had enhancing (weakening) effects when told the device was active compared to a baseline measure. However, while the Stroop task provides a measure of inhibitory control, expectancies about perception, and not inhibition, were manipulated.

Most belief studies, on the other hand, have implemented a quite liberal definition of self-control, encompassing a wide range of actions and mental operations that result from various interacting processes. Breaking down self-control to its underlying functions allows for a better understanding of the effects of expectancies on the constituting facets. This was possible in the current study by combining the emotional antisaccade paradigm with the study of ERPs that reflect specific processes. Hence, capitalizing on the high temporal resolution of EEG, we aimed at identifying the locus of belief influences in the processing stream (between the instruction to suppress an action or not, and the actual inhibition). Cognitive control can be divided into sustained and anticipatory strategies (proactive control) and transient activations of control triggered by stimuli or events (reactive control, Braver, 2012). Based on the timing of the occurrence in the processing stream, the N2/P3-complex can be regarded as reflecting the latter type of control while the CNV can function as a marker of the former (see Vanlessen et al., 2015). The N2 and P3 showed the expected variation with trial type in our data, showing a larger N2 and a smaller P3 amplitude for anti-compared to prosaccades, confirming that the two ERPs captured processes related to the necessity to inhibit. However, groups did not influence these two components differentially. The results for the CNV also showed no difference between the two groups, although a trend was observed towards a lower CNV in the LSC compared to the MSC group. While this effect did not reach significance, it might suggest that expectations rather impact proactive and not reactive control processes as shown by the numerical difference between groups (mean MSC = −0.29 µV/cm^2^; mean LSC = −0.18 µV/cm^2^). However, further research is necessary to investigate a distinction in effects of beliefs or expectations on inhibitory control as well as more complex behavior.

Within the time frame following target onset, we also looked at an attention-related ERP component to determine if beliefs would proactively impact processing of target pictures. This is an informative distinction given recent research showing proactive dampening of attention in a task occasionally demanding inhibitory control (Langford et al., 2016), as is the case in the antisaccade paradigm. Drawing on stronger attentional capture and processing of emotional stimuli (Pourtois et al., 2013), the inclusion of such stimuli in the antisaccade paradigm was aimed to further clarify whether belief modulations would impact the attentional level of processing. Moreover, it also allowed to investigate emotional in addition to response inhibition, as it has previously been shown that individuals can mentally distance themselves from negatively arousing images in a voluntary manner (Kühn, Haggard, & Brass, 2014). However, we did not find effects of emotional valence on attentional processing in the present study, although there was a trend towards a significant saccade x group effect for the P1, stemming from a reversed pattern for the pro– and antisaccade trials in the two groups (higher amplitude for pro–compared to antisaccades in the MSC; lower amplitude for anti –compared to prosaccades in the LSC). To our knowledge, most studies employing emotional targets in the antisaccade task were focused on the effects of emotion in populations with (a tendency for) psychopathology. One study did investigate the effects of emotional targets using this paradigm in a healthy population (Kissler & Keil, 2008). In this study, pro- and antisaccades were presented in separate blocks, and trials with pleasant and unpleasant compared to neutral IAPS pictures elicited more errors in antisaccade trials when gap separated the offset of the cue and the onset of the target. However, a study on social anxiety using avatars with emotional computer-generated facial expressions in a mixed antisaccade task (Wieser, Pauli, & Muhlberger, 2009) and a study on bipolar disorder using emotional facial expressions of actors as target images in separate pro- and antisaccade blocks (García-Blanco, Perea, & Salmerón, 2013) reported no effects of emotion in their healthy control groups. One difference between the studies not finding an emotion effect (including the present one) and the study of Kissler and Keil (2008) is the presence of a gap between cue and target. However, a gap is aimed at rendering the inhibition of erroneous prosaccades more difficult. A lower activity of fixation neurons and increased activity of saccade neurons in the superior colliculus and frontal eye fields during the gap presumably lowers the threshold for triggering a (erroneous) saccade when a target appears (Everling, Dorris, & Munoz, 1998; Mayfrank, Mobashery, Kimmig, & Fischer, 1986; Munoz & Corneil, 1995; Munoz & Everling, 2004). Hence, the gap mostly influences processes specifically related to saccade preparation prior to target-onset, while emotional valence of targets influences processes following its appearance. It is thus likely that stimulus properties, rather than the presence or absence of a gap, contribute to the discrepancy between our study and the one of Kissler and Keil (2008). While the authors controlled the images for brightness, contrast and color spectrum, it is still possible that certain attention-attracting colors were more present in the emotional compared to the neutral stimuli. In the present study, we therefore only used gray-scaled images that were matched on luminosity, contrast and complexity, ruling out additional low-level confounding variables to account for effects of picture valence. For future research, we therefore also suggest to carefully control for such low-level properties to establish valence effects of images in various tasks.

### Placebo/nocebo and beliefs

The central question of this study was whether “pure” belief manipulations can be used to increase or decrease self-control. In other words, can self-control be enhanced or weakened merely by evoking the *belief* that the level of self-control capacity is changed? In this study, we capitalized on the placebo/nocebo effects to induce such beliefs. Placebo/nocebo effects are created through the mechanisms of learning from previous experiences and instructions, although it remains debated whether both are necessary to elicit the effects (Stewart-Williams & Podd, 2004). In an experimental context, the subjective experience of the treatment effects can be evoked by manipulating the experience of participants during a so-called conditioning phase. During conditioning, the experimental task is changed to manipulate the experience of task performance to match the induced expectations by, for instance, adapting difficulty levels so that participants get the impression they perform better or worse (Schwarz & Büchel, 2015) or using false feedback (e.g., in self-efficacy research, Hamburg & Pronk, 2015). However, when both aspects of the placebo context, the instructions and the learning experience, are present, it is not possible to quantify the power of expectations induced with instruction alone. Hence, in our study, we used a placebo/nocebo manipulation consisting of bogus neuro-stimulation without a conditioning phase. Besides circumventing the above-mentioned confound, placebo/nocebo procedures offer the advantage of being less abstract compared to belief manipulations, that often consist of reading a scientific text aimed at changing the participants’ convictions (e.g., Rigoni, Kühn, Gaudino, Sartori, & Brass, 2012).

Our results suggest that placebo/nocebo-inducing instructions alone can evoke a change, both on the subjective and the behavioral level, in line with the study of da Gama and colleagues (2013). In another study, participants were told that specific tone frequencies would impact their functioning on a cognitive control task, followed by a conditioning phase (Schwarz & Büchel, 2015). Results showed that while the subjective feeling of participants about their performance was influenced in accordance to the experimental condition, objective measures were not. The exploratory analysis we performed on the timing of the placebo/nocebo effect in our data might provide a tentative explanation for this discrepancy. This analysis revealed that the interaction effect between trial type and group for accuracy only occurred in the second half of the task. This might suggest that participants compared their estimated level of performance with the expectancy that the brain stimulation made them better (MSC group) or less able (LSC group) to perform the task correctly. Given that most errors are swiftly corrected in the antisaccade task, participants seemed to know when they committed an error and thus generated accurate online feedback about their own performance. In this line of reasoning, participants noticed the discrepancy between the expected and the observed performance throughout the first half of the experiment, and started regulating their behavior to match with the expected performance throughout the second part of the experiment. When the placebo/nocebo manipulation contains a conditioning phase prior to the experimental phase, this initial matching process signals that the performance is aligned with their expectations and that there is no need for regulating performance. However, our study does not allow to allocate the impact of the instructions to one condition or the other (i.e., MSC or LSC group).

Given that our main focus was to establish whether or not instructions alone could elicit a placebo/nocebo effect on measures of self-control, we did not include a control group in which no specific expectancies were elicited. However, it would be interesting for future research to further unravel the mechanisms behind the observed placebo/nocebo effect in our data, and to differentiate the potency of either placebo or nocebo effects on higher-order cognitive functions.

## ACKNOWLEDGMENTS

This research was funded by a grant from the John Templeton Foundation’s Philosophy and Science of Self-Control Project (15462). AS is supported by a postdoctoral mandate from Ghent University (BOF14/PDO/123). The authors have no conflict of interest to declare.

## Footnotes

1 Pilot studies indicated that 15-20% of participants indicate not to believe that the placebo neuro-stimulation resulted in perceived changes in cognitive or behavioral ability to perform tasks. Hence, a large sample size was necessary to ensure around 30 participants per group.

2 We did not include a control group in which no specific expectancies were elicited because our main focus was to establish whether or not instructions alone could elicit a placebo/nocebo effect on measures of self-control.

3 Because of the nature of the task, the latency of the N2 is quite variable. Therefore, a peak analysis instead of a mean area was used for analysis.

