## Supplementary material for "I believe, therefore I am: Effects of self-control beliefs on behavioral and electrophysiological markers of inhibitory and emotional attention control": Table 1

Table 1. Descriptive data of manipulation check questions.

| QUESTION | MEAN | STD. DEVIATION |
|---|---|---|
| tDCS expectancy | 5.17 | 0.92 |
| Think | 4.78 | 0.95 |
| Feel | 4.17 | 1.32 |
| tDCS estimation | 4.36 | 1.26 |
| Attribution | 4.16 | 1.33 |
| Effort | 4.70 | 1.19 |
| Task efficiency | 4.56 | 1.10 |
